# Quasi-Mendelian Paternal Inheritance of mitochondrial DNA: A notorious artifact, or anticipated mtDNA behavior?

**DOI:** 10.1101/660670

**Authors:** Sofia Annis, Zoe Fleischmann, Mark Khrapko, Melissa Franco, Kevin Wasko, Dori Woods, Wolfram S. Kunz, Peter Ellis, Konstantin Khrapko

**Affiliations:** Department of Biology, Northeastern University, Boston, MA, USA; Department of Experimental Epileptology and Cognition Research, University of Bonn, Germany; School of Biosciences, University of Kent, Canterbury, Kent, CT2 7NJ, UK

**Author notes:** Corresponding co-authors,. Equal contribution (Sofia Annis, Zoe Fleischmann).

## Abstract

A recent report by Luo et al (2018) in PNAS (DOI:10.1073/pnas.1810946115) presented evidence of biparental inheritance of mitochondrial DNA. The pattern of inheritance, however, resembled that of a nuclear gene. The authors explained this peculiarity with Mendelian segregation of a faulty gatekeeper gene that permits survival of paternal mtDNA in the oocyte. Three other groups (Vissing, 2019; Lutz-Bonengel and Parson, 2019; Salas et al, 2019), however, posited the observation was an artifact of inheritance of mtDNA nuclear pseudogenes (NUMTs), present in the father’s nuclear genome. We present justification that both interpretations are incorrect, but that the original authors did, in fact, observe biparental inheritance of mtDNA. Our alternative model assumes that because of initially low paternal mtDNA copy number these copies are randomly partitioned into nascent cell lineages. The paternal mtDNA haplotype must have a selective advantage, so ‘seeded’ cells will tend to proceed to fixation of the paternal haplotype in the course of development. We use modeling to emulate the dynamics of paternal genomes and predict their mode of inheritance and distribution in somatic tissue. The resulting offspring is a mosaic of cells that are purely maternal or purely paternal – including in the germline. This mosaicism explains the quasi-Mendelian segregation of the paternal mDNA. Our model is based on known aspects of mtDNA biology and explains all of the experimental observations outlined in Luo et. al., including maternal inheritance of the grand-paternal mtDNA.

## Introduction

A recent report (Luo et al., 2018) presented the long-awaited confirmation of paternal inheritance of mtDNA in humans (Schwartz and Vissing, 2002). Surprisingly, paternal transmission of mtDNA(Luo et al., 2018) follows a bi-modal pattern: about a half of the offspring show fairly uniform paternal/maternal heteroplasmy levels while the rest do not inherit paternal mtDNA at all. This pattern resembles the inheritance of a dominant nuclear gene. The authors explain this pattern as permissive inheritance resulting from a faulty ‘gatekeeper’ gene (Luo et al., 2018). However, three groups (Vissing, 2019), (Lutz-Bonengel and Parson, 2019), (Salas et al., 2019) instead suspect contamination with mtDNA nuclear pseudogenes (NUMTs), a notorious artifact (Hirano et al., 1997).

Based on our vast NUMT experience, we support the authors’ response (Luo et al., 2019), asserting that NUMT artifact is unlikely (***further explanations***: **Notes 1-2**). However, we also demonstrate that the authors’ dominant gatekeeper explanation (Luo et al., 2018) is ***incorrect***, because spermatozoa are functionally diploid (**Note 3**) and thus are equally affected by the faulty gatekeeper. This leaves the quasi-Mendelian inheritance unexplained.

We offer an alternative explanation based on our analysis of the intracellular population dynamics of paternal mtDNA (**Fig.1**): With a defective gatekeeper, a spermatozoid delivers =<100 paternal mtDNA molecules (**Note 4**) to the oocyte. In the beginning, cleavage of the embryo proceeds without mtDNA replication, and mtDNA molecules, including paternal ones, are randomly distributed among the blastomeres. Because the number of paternal molecules is low, some blastomeres are not seeded with paternal mtDNA (**Fig1B**) or lose them due to intracellular genetic drift. Early replication of a mtDNA subpopulation counterbalances genetic drift, with about half of cells getting seeded with paternal mtDNA (**Note 7.2**).

**Figure 1.**
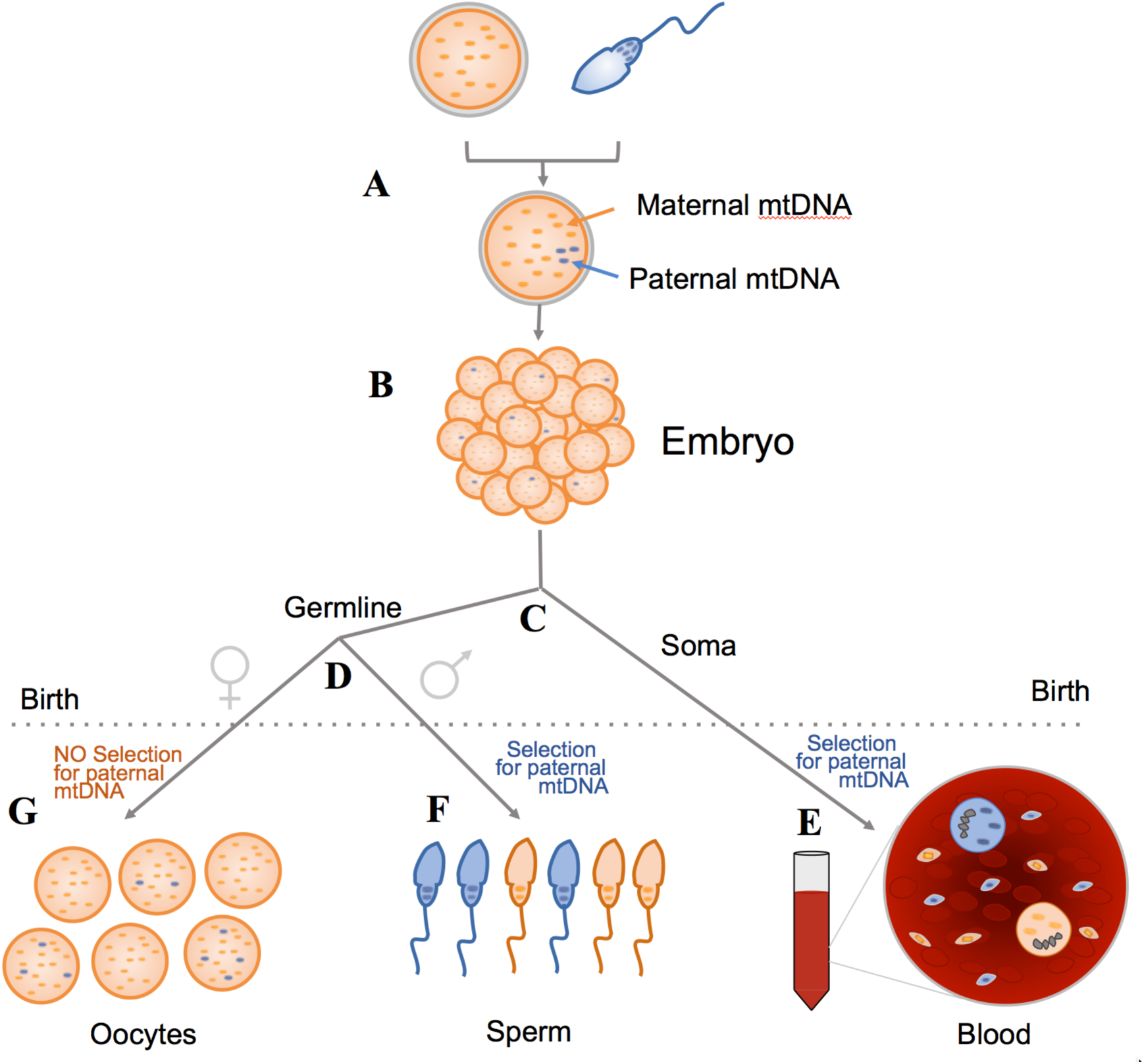
As paternal mtDNA **(A**) is distributed among blastomeres not all of them receive a copy (**B**). Later germline is parted from soma (**C**) and then the sex of the germline is set (**D**). After **birth** (dotted line), somatic and male germline cells exert *selection* in favor of the paternal mtDNA haplotype, so that somatic cells and spermatozoa end up as a paternal/maternal mtDNA mosaic. This explains somatic heteroplasmy and quasi-Mendelian inheritance, respectively. In the *female* germline, there is no selection for the paternal haplotype (8), so oocytes keep low and variable proportions of paternal mtDNA (**G), (Note 7.3.2**)

Because paternal inheritance includes *reproducible,* ∼1000-x enrichment of paternal mtDNA, paternal mtDNA haplotype *must* have a selective advantage over maternal haplotype (**Note 5**). Competition between mtDNA haplotypes in somatic tissues is well-documented (Jenuth et al., 1997), and can be sufficiently strong to drive the *enrichment* of the paternal haplotype to ∼100% in the seeded cell lineages (**Note 6**). Non-seeded lineages naturally remain 100% maternal, so a mosaic of cells with mostly paternal and pure maternal mtDNA is created (**Fig.1E**). In mosaic tissue, the heteroplasmy level is equal to the proportion of paternally-derived cells, or seeding density i.e., the fraction somatic cell lineages that received and kept paternal mtDNA during embryogenesis.

Similarly, not all germline lineages are seeded with paternal mtDNA, which ultimately results in a mosaic of paternally- and maternally-derived spermatozoa (**Fig.1F**). This explains the quasi-Mendelian bi-modal inheritance, with a transmission rate equal to the seeding density of germline lineages. Maternal germline follows a different fate (**Fig.1G, Note 7.3.2**)

Intuitively, seeding density should depend on the initial number of paternal mtDNA, the timing of mtDNA replication initiation, and the dynamics of the intracellular population of mtDNA molecules. To investigate the combined effect of these factors quantitatively, we performed *numerical simulations of mtDNA behavior* (**Note 7.2**). Simulations predict intermediate seeding densities and heteroplasmy levels in somatic and germ cells (**Note 7.3**), in agreement with the observations (**Note 7.1**).

In conclusion, despite apparent inconsistencies and some missing quality control items (**Note 10**), Luo et al. likely describe genuine cases of paternal inheritance, and seemingly impossible inheritance patterns are compatible with known mtDNA biology.

## Methods and Results

### Note 1. NUMT contamination: an unlikely possibility

While this letter was in preparation, a recent letter (Lutz-Bonengel and Parson, 2019) and author’s response (Luo et al., 2019) treated the problem of NUMT contamination in some detail. Because some of us have been involved in the research on NUMT effects in mtDNA mutational analysis since the issue was first discovered in early nineties by Gengxi Hu in the Bill Thilly’s laboratory at MIT (Khrapko et al., 1994), we would like to add our perspective to the NUMT contamination problem.

#### Note 1.1 mtDNA-specific damage boosts the perceived NUMT fraction

In our experience, one of the causes of apparent NUMT contamination is mitochondrion-specific DNA damage. Specific damage of mtDNA was the cause of NUMT contamination in the case when this problem was first identified (Hirano et al., 1997). While specific damage of mtDNA does decrease the apparent number of mtDNA copies in a sample and in severe cases may result in parity of mitochondrial and nuclear DNA or even the prevalence of nuclear DNA, this is not typical in DNA isolated from fresh or frozen blood using modern laboratory practices – it’s more typical of partially degraded autopsy samples, etc.

##### Suggested test for mtDNA damage

The presence of mtDNA damage can be retrospectively tested on the existing samples of purified DNA from blood samples: PCR should be performed using much shorter PCR fragments. The effect of damage dwindles dramatically with shorter fragment length, so if NUMT contamination is involved because of mtDNA-specific damage, shorter PCR will result in a sharp decrease of the paternal haplotype fraction.

Another factor could be cell lysis prior to DNA extraction. If the sample is not stored carefully, then the plasma membrane might be damaged, leading to loss of cytoplasm. Then, when the cells are centrifuged to extract the DNA, only nuclei may be pelleted, while the mitochondria may be lost in the supernatant resulting in enrichment for nuclear NUMTs. This possibility emphasizes the need for other tissue measurements (such as buccal cells). We note however, that chances are that the paternal haplotype is enriched in blood but not in the buccal cells. Indeed, it is known that NZB/Balb selection differ dramatically between different tissues (from positive to negative). Similarly, in the well-established case of paternally inherited mtDNA (Schwartz and Vissing, 2002) the paternal haplotype strongly showed up in muscle, but was completely absent in blood. So, ideally, the confirmatory tests should be performed in blood.

#### Note 1.2 NUMT must be full length, tandem repeat of a perfect mtDNA copy

The scheme in (**Fig.2**) shows that in the case of linear mtDNA copy inserted into nuclear genome, one’s ability to perform the full length PCR with either of two different pairs of primers (as reported in materials and methods, (Luo et al., 2018)), requires that the flanks of the pseudogene sequence in the genome are also mtDNA sequences continuously extending the sequence of the central copy, as if this was a circular mtDNA molecule. Biologically such a situation most likely can arise by tandem multiplication of the unaltered mtDNA sequence in the nuclear genome.

(Luo et al., 2018) amplified mtDNA using two separate pairs of primers (**A** and **B**) that each generated nearly full-length products (green and orange lines). If the observed paternal haplotypes were indeed NUMT contaminants, the NUMT would have to be a perfect tandem repeat that encompasses the forward and reverse priming sites for both primer combinations

While existence of full length tandem repeat of unaltered copies of mtDNA in the nuclear genome cannot be completely excluded, such an arrangement has not been observed in sequenced human genomes or other genomes. Even simply large mtDNA fragments are very rare in the human nuclear genome. There are a few cases of almost full genome NUMTS – one recently detected in a cancer tissue (Yuan et al., 2017), one in a columbine monkey (Wang et al., 2015), but all of them contain at least some deletion. Moreover, NUMTs appear to be rather stable in the nuclear genomes of various species. We have observed a 5kb NUMT which did not change (other than acquired point mutations) for at least 8 million years while residing in Human, Chimp and Gorilla genomes (Popadin et al., 2017). The columbine monkey NUMTs also showed no structural change since their divergence about 4 million years ago (Roos et al., 2011). This implies that the lack of tandemly repeated perfect mtDNA copies is likely not a result of disappearance of such structures with time but rather their failure to arise in the first place.

In conclusion, while the presence of perfect multicopy NUMTs in the families described in Luo 2018, cannot fully excluded, this appears to be a very unlikely possibility. If such NUMT arrays can be proven to exist, that would be, in our opinion, a discovery similar in scientific value to the confirmation of the paternal transmission case.

### Note 2. Tests for NUMT contamination: platelet and nuclei analysis

The NUMT hypothesis can be most easily tested by comparing heteroplasmy levels in tissues with different mtDNA/nDNA ratio. For example, in case of Luo et al., in blood samples, analysis the purified platelet fraction would help. Platelets contain no nuclear DNA, so if NUMT hypothesis is correct, the proportion of the paternal genotype should be drastically reduced when re-measured in purified platelets. Platelets should be easy to isolate even from frozen archived blood samples, so this is hopefully something that can be done even if new sampling is not possible.

In addition, it would be important do the converse analysis by purifying nuclei from a blood sample using hypotonic lysis. That should eliminate most mtDNA and enrich NUMTs. Moreover, the degree of enrichment of nuclear DNA can be directly checked using PCR to test for constitutive NUMT sequences.

#### The original case of paternal inheritance is not a NUMT

Of note, the first detected case of paternal inheritance (Schwartz and Vissing, 2002), did pass a similar NUMT test: the fraction of paternal mtDNA was high in muscle, a tissue rich in mtDNA, and undetectable in blood, where relative mtDNA content is low, i.e. the extreme reverse of what NUMT hypothesis would have predicted. Also as we explored the presence of mtDNA recombinants (essentially products of reciprocal repair), we extensively tested the DNA from the patient (both blood and muscle) at different PCR fragment lengths, and detected no difference in the maternal/paternal haplotype ratio (Kraytsberg et al., 2004), arguing against differential damage possibility (**Note. 1**). Of note, because the original case of paternal inheritance (Schwartz and Vissing, 2002) passed both tests, the NUMT hypothesis at least is not generally applicable (**Note 4**).

### Note 3: Functional diploidy of spermatozoa: Mendelian segregation of a gatekeeper gene is not a valid explanation of bi-modal inheritance

Luo et al (2018) propose that the quasi-Mendelian inheritance pattern exhibited among the offspring of heteroplasmic fathers that transmit their mitochondria is due to the underlying segregation of an autosomal dominant gatekeeper gene mutation that promotes paternal mitochondrial transmission. Under this hypothesis, the gatekeeper gene must function cell-autonomously after meiosis, i.e. only those sperm cells that inherit the mutant allele pass on their mitochondria. This contradicts the known biology of germline development. A fundamentally conserved aspect of spermatogenesis throughout the animal kingdom is that cytokinesis during spermatogenesis remains incomplete during the premeiotic and meiotic divisions. This means that all the sister cells arising from the same germline stem cell remain linked by cytoplasmic bridges (BURGOS and FAWCETT, 1955),(FAWCETT et al., 1959). The bridges are large, up to 3 microns in diameter (Weber and Russell, 1987), i.e. sufficient size and scale to enable sharing of cytoplasmic contents between cells (Braun et al., 1989). This sharing is an active process, in which mRNAs are trafficked between cells via specific binding proteins and shuttling of transcripts within a dedicated organelle known as the chromatoid body. (Morales et al., 2002), (Ventela et al., 2003). In addition to the facilitated transport of mRNAs within the cytoplasm, both rough and smooth endoplasmic reticulum are continuous across the intercellular bridges, allowing sharing of protein products between cells (Clermont and Rambourg, 1978), figure 18. Consequently, sperm are functionally diploid, not haploid.

Could it be possible for the gatekeeper gene to avoid transcript sharing and act in a cell-autonomous manner? While theoretically possible, this would be virtually unprecedented since only two examples of non-shared genes are known in mice ((Zheng et al., 2001), (Veron et al., 2009)) and none in humans. However, even a non-shared gatekeeper gene cannot explain the observed inheritance pattern in the families studied by Luo et al. Whole mitochondria have been observed within the intracellular bridges by electron microscopy (Dym and FAWCETT, 1971) figures 19 and 20), implying that mitochondria are also freely exchanged between sister cells. Thus, the mutant gatekeeper gene product would not only have to escape sharing across the bridges, but also modify the mitochondria in such a way that they are no longer shared across the bridges. We judge this to be extremely implausible. Consequently, any gatekeeper mutation, whether dominant or recessive, should affect all sperm cells equally regardless of which alleles they happen to inherit during meiosis.

### Note 4: How many paternal mtDNA molecules are delivered (and survive) by the spermatozoid into the oocyte in the cases of paternal inheritance?

The number of mtDNA in a sperm is controversial, and estimates vary from ∼100 to essentially none. This variation may be due to the self-destructive potential for elimination of mtDNA in spermatozoa, as described (Luo et al., 2013). This may be a premature artificial *in vitro* phenomenon, and in vivo sperm in fact may enter the oocyte with its mtDNA still intact. We therefore assume that once the elimination of sperm mtDNA is disabled, we may expect that about 100 paternal mtDNA molecules will survive in the fertilized oocyte. It might be even less, since the mid sperm tail, where the mitochondria are localized does not penetrate the egg. Nevertheless, it’s probably safe to assume that a substantial fraction of sperm mitochondria (and, hence, mtDNA) are injected into the oocyte, since (Rojansky et al., 2016) conservatively observed ∼40 paternal mitochondrial objects in the fertilized oocyte at 36hrs (two cell embryo). This count is conservative in that overlapping mitochondrion images were counted as one, and also in that counts were taken at 2-cell stage, at which it has been shown that the disaggregation of paternal mitochondria is far from complete (Luo et al., 2013), so many individual mitochondrial objects probably still stick to each other and are counted as one. This interpretation is supported by the highly uneven luminosity of the objects, implying that some of them are aggregates (see Figure 2 from (Luo et al., 2013)) and the number of objects increases at least until the morula stage. Importantly, when sperm mtDNA does make it into the oocyte, it can be identified and traced all the way to multiple tissues of the newborn (Luo et al., 2013).

**Figure 2.**
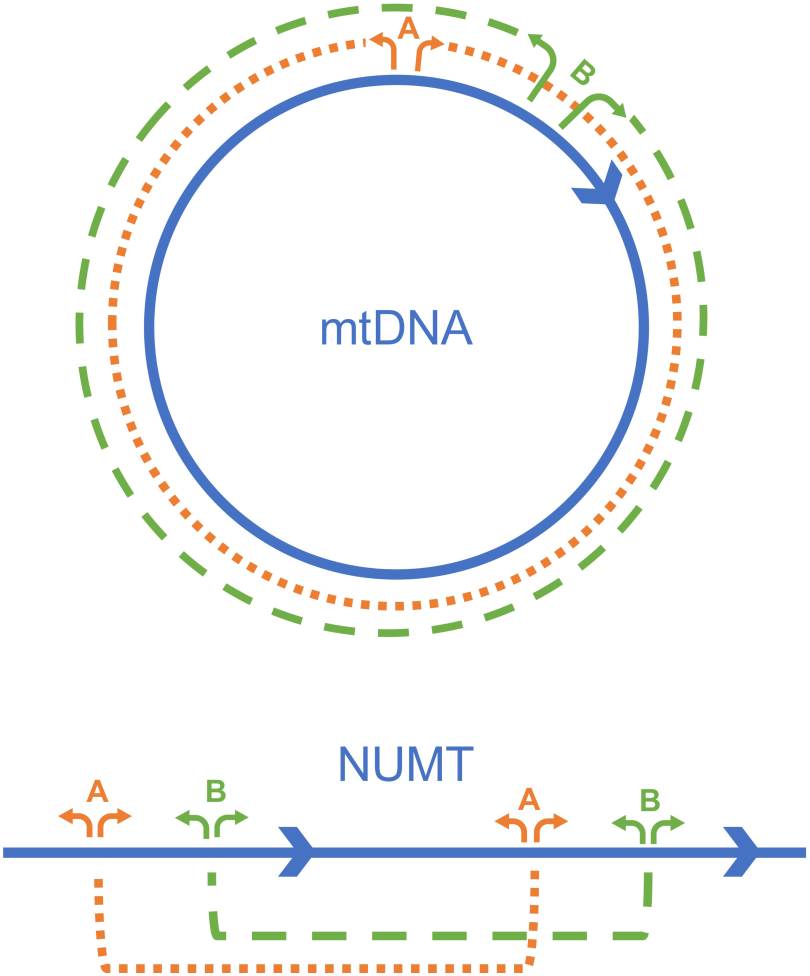

We note that paternal mtDNA copies delivered by the spermatozoid is one of the inexact parameters of our model, and lower than we assumed number of delivered copies may be compensated by earlier than we assumed onset of mtDNA replication and smaller replicating subpopulation (Note 7.3.3).

### Note 5: *Reproducible* paternal mtDNA transmission requires *selection* of the paternal haplotype

#### Note 5.1 Paternal expansion is reproducible

Paternal transmission described by (Luo et al., 2018) is **reproducible**, meaning that it is repeatedly observed within a family. In each of the three families described in (Luo et al., 2018), paternal transmission must have happened more than once. This may not be immediately obvious from the tree itself: there is only one case (family C) where more than one child clearly inherits paternal mtDNA (CIII6&7). Nevertheless, paternal transmission should have happened more than once in all three families because in each family, the path of paternally inherited mtDNA starts with a heteroplasmic man, who already carries two highly divergent haplotypes. Even if we do not consider the great-grand parents, whose DNA was not available and genotypes are uncertain (despite the author’s attempt to infer them), the very existence of the hetero-haplotypic person implies that at least one additional paternal inheritance event have happened sometime in the past to initially create this unusual distant heteroplasmy of two haplotypes. This means that in all three families, paternal transmission was indeed **reproducible**.

The reproducibility of paternal transmission is conceptually very important. Paternal inheritance of mtDNA in cases reported by Luo et al. includes reproducible = ∼1000-x enrichment of paternal mtDNA, from ∼0.05% in the egg (i.e., ∼100 molecules among 200,000 maternal molecules) to ∼50% heteroplasmy in blood. While such enormous enrichment apparently can happen due to random drift alone (next paragraph **Note 5.2**.), such events are quite rare and the probability that two such events happen within the same family is very small, so a selective process is the only reasonable explanation of *reproducible* paternal transmission of paternal mtDNA in a family.

#### Note 5.2 Random enrichment of paternal mtDNA

The efficient enrichment of a rare mtDNA variant in one generation can happen, though rarely, via a random selection-free process; this is basically a manifestation of the mtDNA germline bottleneck phenomenon, which in turn is probably based on intracellular random genetic drift. Perhaps the most dramatic example of random enrichment is the expansion, in tumors, of random somatic mtDNA mutations, including fully synonymous ones, which excludes the possibility that any selection is involved in the process of their enrichment. These mutations appear initially in a single mtDNA molecule and remarkably make it to 100% of the entire tumor (Polyak et al., 1998). We have previously shown that this process can be realistically modeled as random intracellular genetic drift (Coller et al., 2001). As far as the germline is concerned, the abrupt enrichment of one genotype over the other have been recognized as the mtDNA bottleneck effect since a seminal work in heteroplasmic cows (Hauswirth and Laipis, 1982). More recently, extensive population studies on human mother/offspring pairs showed a significant number of cases where a genotype that was not detectable in the mother showed up as significant heteroplasmy in the child (Rebolledo-Jaramillo et al., 2014), (Li et al., 2016). This implies that a rare genotype, perhaps a nascent mutant molecule, in some cases may enjoy a dramatic enrichment among germline mitochondria in normal humans within a one generation time frame.

If paternal mtDNA that has leaked into the oocyte due to defective gatekeeper happens to appear in the role of such a randomly enriched sporadic genotype (and there is no reason why it can’t be in this role), the paternal genotype may overcome the maternal mtDNA and result in paternal inheritance. In fact, this is the same path that any neutral nascent mutation that eventually gets fixed in an organism has to take. And because new neutral mutations do get fixed from time to time, we know that this path indeed exists and is quite efficient on the evolutionary time scale. This implies that reproducible selection is not necessarily needed for paternal inheritance, and thus the only condition of paternal inheritance is the defective gatekeeper. While the coincidence of a lack of paternal mtDNA elimination and random enrichment is still a very rare event, this scenario explains how any haplotype (not necessarily one enjoying reproducible selective advantage) can be potentially paternally inherited.

### Note 6 Haplotype selection of mtDNA – a known process, sufficient to explain paternal mtDNA enrichment observed in paternal inheritance cases

Strong sequence-dependent selection is a natural property of mtDNA. The most studied example is the heteroplasmic NZB/BALB mouse, constructed by cytoplasmic fusion of between mice with highly divergent haplotypes (Jenuth et al., 1997). Relative proliferative fitness of the NZB haplotype is as high as 1.16 per duplication, depending on the tissue type (Battersby and Shoubridge, 2001). This means that the proportion of the NZB haplotype increases 1.16 times per mtDNA duplication. Interestingly, this selective advantage mechanistically may be related to slower turnover in the NZB haplotype rather than faster replication (Battersby and Shoubridge, 2001). Note that with fitness of 1.16 per duplication, it would take fewer than 50 duplications to achieve a ∼1000-fold enrichment that is needed to equalize maternal and paternal mtDNA when the latter starts from 0.05% (indeed, 1.16^50^ ∼1,700). It is not unrealistic to expect hundreds of cell duplications in the somatic tissue cell lineage, which gives the lineage more than enough cell duplications to achieve the necessary ∼1000x enrichment of the paternal haplotype. Also, NZB/BALB enrichment was essentially a randomly chosen pair of haplotypes; it is easy to imagine that other combinations of haplotypes may result in even higher relative fitness. Furthermore, (Luo et al., 2018) cases are not average ones, but were actually selected for *detectability* among large set of samples. Thus the haplotype combinations in these samples are expected to be the ones that display appropriate relative fitness and end up as a detectable heteroplasmy/paternal inheritance.

### Note 7: A detailed model of paternal mtDNA transmission

We devised a numerical model that simulates the intracellular behavior of individual mitochondria in proliferating cell lineages. In this model, intracellular populations of mtDNA molecules are represented by virtual cells, i.e., lists of sequences (some of which are identified as paternal and some as maternal). Sequences can be duplicated (representing mtDNA replication) and removed (representing cell division and/or mtDNA turnover) in any order and proportion. This model allows for adjustments in the number of mitochondria per cell, the size of the replicating subpopulation, the proportion of molecules of each haplotype, their relative proliferative fitness, and the number of cell divisions. Everything is done biologically reasonably, e.g. to increase cell size, we duplicate more sequences (randomly chosen) than we remove, and thus create a positive balance in the number of sequences per virtual cell. The model is composed of several sequential blocks with different parameters representing the different developmental stages. The numbers of germline cell divisions were chosen according to classical work (Drost and Lee, 1995). The algorithm is written in Python and is conceptually based on our original numerical simulations of mtDNA intracellular dynamics in cancer lineages (Coller et al., 2001).

This model is capable of predicting the observables of the Luo et al 2018 study: the transmission rates and the paternal heteroplasmy levels of somatic tissues in the offspring. As we show below, the predictions are in a reasonable agreement with the observations. This does not necessarily mean that processes are in the foundation of our model are necessarily operating in vivo. The goal of our modeling was to show that Luo’s observations, however enigmatic they are, are compatible with a biologically plausible scenario. This goal has been accomplished in full.

#### Note 7.1: The observables to predict: Transmission rates of paternal mtDNA and heteroplasmy levels in the offspring

First, we need to determine what are the observables that our model is expected to predict. For the purpose of this discussion we concern the transmission rate of paternal mtDNA among families that have already shown paternal transmission (not in the general population, where it is obviously very low). In particular, we will use two rates: a) the transmission rate from fathers already known to be transmitters of mtDNA to their offspring, and b) from mothers carrying paternally transmitted mtDNA to their offspring. Note that when counting transmission events we need to exclude any parent/offspring transmissions to or directly upstream of the probands, i.e. individuals in whom paternal transmission has been initially discovered during search. The probands are positive for paternal transmission by the anthropic principle: if they did not show paternal transmission, the families would have not been included in the analysis. With that in mind there are 6 such transmissions in all there families where offspring were analyzed (to AII1, 2, 3; to AIII7; to BIII1, and to CIII7), of which 3 (to AII1 and AII3 and to CIII7) are positive (Luo et al., 2018), i.e. observed transmission rate is **∼50%**.

As far as the rate of maternal transmission of a paternally acquired haplotype is concerned, there are 3 independent events of this type: transmission to individuals AIV1 and AIV3 (AIV2 is omitted as proband), and CIV1 (Luo et al., 2018). All these transmissions are positive, so transmission rate here is 3/3 – **100%**. Of course, the number of events is extremely low, so this number perhaps is a rough approximation only. It looks like transmission rate along maternal line is higher than along the paternal one. Reassuringly, our detailed model (**Note 7.5**) does predict a higher transmission rate in females than in males (this is related to a larger cell size in the female germline).

The average heteroplasmy level of paternally transmitted mtDNA (positive individuals only) is 0.48 ± 0.16. Note that estimating the heteroplasmy levels does not require exclusion of the probands and their direct ancestors because being positive carries very little information about the actual heteroplasmy levels (we are interested in the heteroplasmy levels in the positive individuals only). The average heteroplasmy level of maternally transmitted paternal mtDNA is 0.33 ± 0.15

#### Note 7.2 Model step-by-step: Justification of the scenario and of the parameters used

##### Note 7.2.1 Block 1:Fertilization and cleavage of the embryo prior to the re-start of mtDNA replication

We assumed that ∼100 paternal mitochondria (Note 4, Note 7.3.3) were randomly distributed among 16 blastomeres in the morula. The morula blastomere is expected to contain ∼12,000 (200,000/16) total mtDNA molecules, of which there should be, on average, 6 molecules of paternal mtDNA (100/16=6). So a cell lineage at this stage was simulated by randomly distributing an average of 6 paternal genomes (Poisson-distributed) among virtual blastomeres, i.e., lists 12,000 sequences long.

###### Justification

After entering the oocyte, paternal mtDNA temporarily persists as a cloud of mitochondria, which gradually dissipates and gets uniformly dispersed around the embryo by the morula stage, as recorded in studies where paternal mitochondria were stained with specific fluorescent tags (Rojansky et al., 2016), (Luo et al., 2013). The initial clustering of paternal mitochondria might naturally result in an excess of lineages lacking paternal mtDNA (and others with overcrowded mtDNA), however with the data at hand there is no way we can reliably model this deviation, so we do not take it into consideration. Our estimates are conservative with respect to the expected rate of paternal transmission and can overestimate the level of paternal heteroplasmy for this reason.

##### Note 7.2.2. Block 2: Onset of mtDNA replication: the timing of the onset and the size of the replicating subpopulation

We assumed that mtDNA replication resumes within the mtDNA subpopulations containing 1000 molecules (∼10%) within the morula blastomeres, and that all paternal mtDNA molecules are replicating. 10% for the relative size of the replicating pool was based on our preliminary estimates in PolG oocytes as discussed in the ‘Justification’ below (see also ‘Note of caution’ 7.3.3). The virtual replicating subpopulation was uniformly duplicated 3 times while the entire population of mtDNA is *equally but randomly* halved after each duplication to emulate the ongoing cleavage (thus replicating population oscillated between 1000 and 2000). *Equally but randomly* means that the total mitochondrial population is divided in half (reflecting approximately equal distribution of mtDNA among blastomeres), but the few paternal copies are distributed randomly (and as a being a small number, are subject to much larger relative variance). We assumed that the rest of the mtDNA population either undergoes mitophagy (according to the (Spikings et al., 2007) data) or is diluted out by the newly replicating mtDNA. The completion of this block mimics the implantation stage with ∼128 cells, each now dominated by the replicating mtDNA population, 1000-molecules strong. We then performed 2 cycles to emulate post-implantation proliferation of embryonic cells prior to the commitment to the germline at CS5. Note that during this period the average number of mtDNA molecules per cell is 1500 ((1000+2000)/2=1500), which is in keeping with current estimates (Floros et al., 2018)

###### Justification

The timing and the extent of mtDNA replication reinitiation in the embryo is a critical parameter in our model, so we will spend extra time on the justification of this parameter choice. In particular, it is likely (see below) that mtDNA replication in the embryo starts ***earlier*** than conventionally assumed, but is initially limited to a small subpopulation of mtDNA molecules. Paternal mtDNA is likely to be a part of this replicating subpopulation, which means its frequency in the embryo should get a boost.

In support of the idea of partial replication, note that replication machinery is being gradually rebuild in the preimplantation embryo, and most likely will start to work at partial capacity. Independent studies indicate that mtDNA replication may be starting as early as in the morula (16-cell embryo), because this is the point where a significant amount of mtDNA-replication-related proteins (PolG and TFAM) have been already produced *de novo* in significant amounts (Bowles et al., 2007) (Fig.1 therein). In addition, the intrinsic ability of the early embryo to replenish its mtDNA has been demonstrated in experiments on the depletion of oocyte mtDNA in cattle (Chiaratti et al., 2010). Depleted mtDNA have been fully replenished by as early as the blastocyst stage.

A more dramatic observation is that apparently mtDNA in the early embryo is degraded to a significant extent and then is replenished by replication of a limited stock. According to (Spikings et al., 2007) (Fig 3 therein), total mtDNA copy number in the embryo is sharply reduced in the 8-cell embryo, and then rebounds to pre-cleavage levels by the expanded blastocyst stage. These observations may seem to contradict other studies that reported relatively constant amount of mtDNA in the early embryo, such as (Cao et al., 2007). We note however, that mtDNA content of the embryo is typically assessed by real time PCR that employs a very short PCR fragment, about 100nt long (Cao et al., 2007). In contrast, (Spikings et al., 2007) used a 1000 bp PCR fragment, which is far more sensitive to the acute mtDNA degradation, which should take place in the embryo after activation of autophagy in the 4-cell embryo (Tsukamoto et al., 2008). Note that mtDNA that has just started to be degraded may not have had enough time to have been cut to 100nt fragments, so regular 100-bp real time PCR may not be sensitive enough to the initial stages of degradation, which is later masked by subsequent replenishment of mtDNA in the embryo by *de novo* replication.

Of note, the conventional idea that there is no mtDNA replication until implantation stage is mostly based on the observation that the total amount of mtDNA in the preimplantation embryo is roughly constant. However the above arguments imply that the apparent stability of the DNA content may be the result either of initial replication of a small subset of mtDNA which thus is not expected to significantly affect the total mtDNA amount and this remain undetected, or reflect a balance between replication and degradation.

The observation that mtDNA is degraded and then replenished in the early embryo is further corroborated by the report of intense segregation of mtDNA haplotypes in the chimeric preimplantation embryo combined from two oocytes containing heterologous mtDNA (Lee et al., 2012). This effect is very difficult to explain without assuming a dramatic degradation/replenishment of the mtDNA population. mtDNA degradation in the early embryo actually makes a lot of practical sense: because there is no mitophagy in the oocyte (Boudoures et al., 2017), apparently much of the mtDNA in the oocyte has not been renewed for many years of the oocyte’s lifetime and perhaps is significantly damaged and likely to fail replication.

In corroboration of this view, we independently observed that only a small subpopulation of mtDNA in the mature oocyte (**∼10%**) have been actively replicated, and that mtDNA molecules from this subpopulation are preferentially inherited in the next generation (Annis et al., in preparation).

Early replication of a subpopulation of mtDNA significantly affects our model as long as paternal mtDNA specifically is a part of this replicating subpopulation. The reason to expect this is that unlike a majority of mtDNA of the oocyte, paternal mtDNA has been recently replicated as a part of proliferation in the spermatogonia and thus is expected to be free of replication-impeding damage. Moreover, sperm mtDNA is pre-loaded with PolG and TFAM (St John et al., 2007) (Figure 3 therein), putting it in the position to start replicating ahead of the rest of oocyte mtDNA, which needs the de-novo protein synthesis to provide PolG and other factors essential for mtDNA replication.

##### Note 7.2.3 Block 3: Germline/soma specification to birth

At CS5, after 9 embryonic cell divisions, the germline is specified as a separate group of cells, Primordial Germ Cells (PGCs), which have distinctly larger size than soma-to-be cells. In keeping with this milestone, from this point simulations were carried for 7 duplications along two paths representing the germline and the somatic lineage. The separate paths reflect the different average size of the primordial germ cells (PGCs, 1500 mtDNA/cell) and somatic cell lineages (500 mtDNA/cell) (Cao et al., 2007), and the size of virtual cells was adjusted accordingly. We used the same approach as in previous blocks: 2-fold oscillation around the average cell size, 7 cell duplications were performed in this regimen. After 7 cell duplications, at the point corresponding to CS17-18, the germline is subject to sex differentiation resulting in differently sized male and female germline cells (∼1750mtDNA/cell on average for female PGCs, ∼1000 mtDNA per cell for male PGCs (Floros et al., 2018)). After that, the germline is carried through 13 more cycles until birth (as before, two-fold oscillation around average cell size).

While the germline was carried through 7+13 cell divisions, somatic cells were carried through a different path with smaller virtual cells. An important question is how many generations does the somatic lineage go through prior to birth. We found no definite answer in the literature and apparently the actual number of duplications can vary significantly even within the same cell type (Reizel et al., 2012). We thus performed simulations with a range of number of duplications: 20, 30, 40, which essentially covers the entire range of plausible possibilities. These differences in duplication number resulted in only moderate deviations of the anticipated heteroplasmy levels (∼20%), so we have chosen to present the entire range in the results table (table 1). The whole range is sufficiently narrow to support our conclusions.

**Table 1:** Predictions from virtual cell simulations compared to observed inheritance rate or heteroplasmy level.

|  | **Predicted:**<br>fraction of<br>lineages seeded<br>with paternal<br>mtDNA | **Observed:**<br>inheritance rate or<br>heteroplasmy level |
| --- | --- | --- |
| Male germline | **0.66** | **0.5**<br>**(inheritance rate)** |
| Somatic cells<br>(male inheritance) | **0.28-0.44\*** | **average: 0.48**<br>**(heteroplasmy level)** |
| Female germline | **0.76** | **1**<br>**(inheritance rate)** |
| Somatic cells<br>(female<br>inheritance) | **[0-1]**<br>**distribution**<br>**see Fig. S3B** | **0.24\*\*, 0.57**<br>**(heteroplasmy level)** |

\* range reflects the models assuming 40 to 20 duplications of the somatic cell lineages
\*\* average of 3 siblings, see Note 7.3.2 for explanation)

##### 7.2.4 Block 4: Post-natal mtDNA behavior

Male germline and somatic cell lineages were simulated with selective pressure in favor of the paternal haplotype. To emulate selective pressure, in each duplication, every fifth randomly selected paternal mtDNA molecule was replicated twice, to represent the ∼20% higher fitness of the paternal haplotype. Mechanistically this bias most likely results from slower turnover of the fit haplotype, not from the faster replication (Battersby and Shoubridge, 2001), but this does not matter as far as the simulation is concerned. The simulations with selection pertain only to the somatic lineages and post-puberty male germline lineages; female germline is assumed not to be subject to selection (Jenuth et al., 1997).

Post-natal behavior of mtDNA in the female germline was simulated by proportionally increasing the mtDNA populations of female PGCs (obtained in the process (3) above) to 200,000 genomes – reflecting the oocyte growth process. For example, a PGC containing 2 paternal mtDNA among 1000 mtDNA was expanded to a 200,000 mtDNA oocyte with 400 paternal mtDNA molecules. Because the replicating subpopulation of mtDNA is only 10%, 40 out of 400 paternal genomes were assigned to this subpopulation. The resulting virtual oocyte was treated as described above for the oocytes fertilized with sperm with paternally inherited mtDNA. The difference is that in this case mtDNA from next generation spermatozoid contributed no paternal mtDNA (because gatekeeper was intact and paternal mtDNA was destroyed); instead, grand-paternal DNA has been carried over from preceding generation through the female germline.

#### Note 7.3 Simulation Results

The most important conclusion from our simulations is that, first, fairly simple and biologically reasonable assumptions (like early onset of proliferation of a small subset of mtDNA) are sufficient to explain the necessary boost of the initially infinitesimal fraction of the paternal mtDNA to the observed high heteroplasmy levels. Second, it explains the observed differences/similarities between heteroplasmy levels and transmission rates in males and females and between germline and somatic cells (which in our model are linked to each other in a complex but sensible way). It should be noted that the model’s reaction to parameter changes is not trivial – this is a system of checks and balances. For example, moving the onset of mtDNA proliferation into an earlier developmental stage helps to explain the remarkable gain in paternal mtDNA frequency and high paternal transmission rate, but going too far results in a too-high transmission rate and heteroplasmy levels in *maternal* transmission of paternal haplotype. The ability of the model with a biologically reasonable set of parameters to satisfy complex and often conflicting observations gives us additional confidence in our conclusions. The results are described in more detail below.

**Figure S2.**
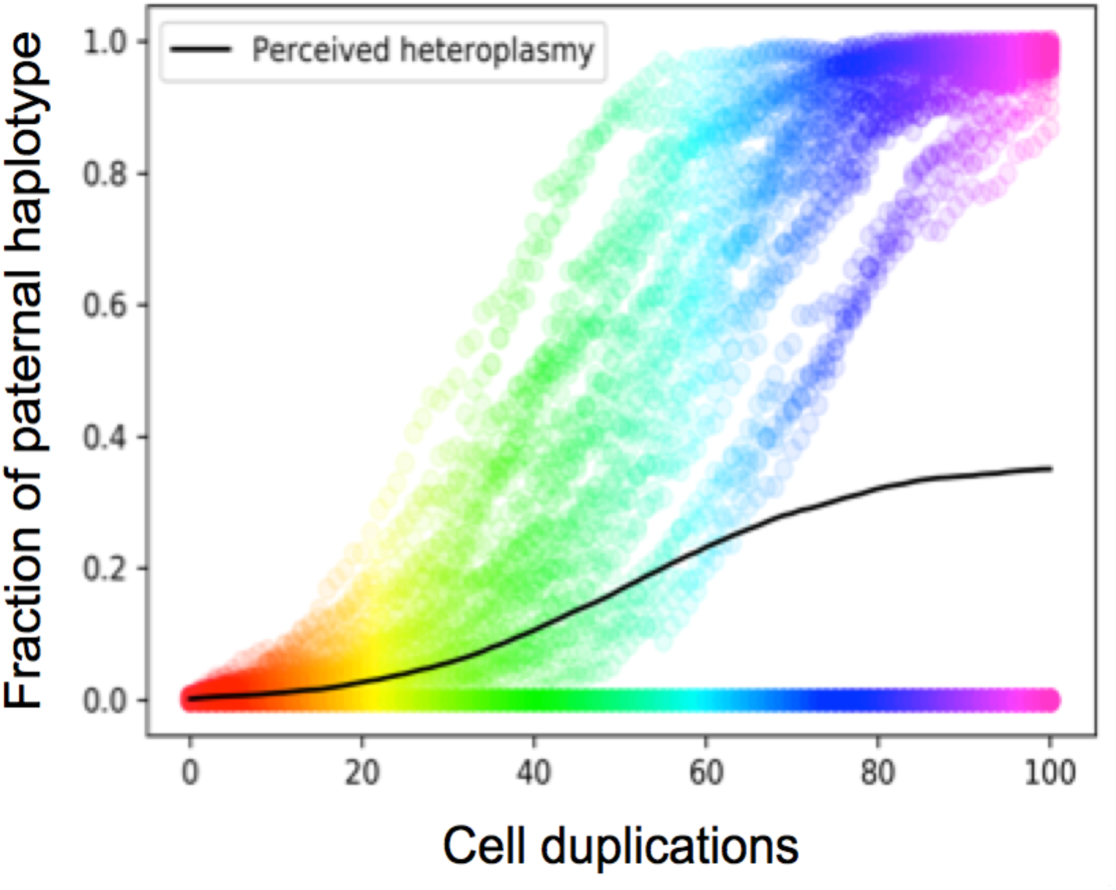
Population dynamics of paternal haplotypes under positive selection: preservation of seeded cell lineages. 100 cells (500 genomes per cell) were randomly seeded with 0.5 paternal genomes per cell on average (which resulted in an initial seeding density of ∼0.39). The cells were then carried through 100 cell duplications with the 1.2 relative fitness assigned to paternal molecules. The graph shows gradual expansion of paternal haplotype in the lineages. Importantly, the asymptotic perceived heteroplasmy, i.e., the average across all cells (∼0.36) appears essentially equal to the initial seeding density of the paternal haplotype (0.39), implying that there is essentially no additional loss of seeded lineages under positive selection regime at seeding densities used in our simulations. Colors reflect the progression of simulations.

##### Note 7.3.1 Rate of paternal inheritance and levels of somatic heteroplasmy of paternal mtDNA in the offspring

The main output of our modeling is the fraction of cell lineages seeded by the paternal haplotype by the time of birth, at which time haplotype selection turns on (Jenuth et al., 1997). According to post-natal simulations (with selection), this transition effectively prevents any further loss of the paternal haplotype from cell lineages. This is because the intracellular genetic drift that had been primarily responsible for haplotype loss slows down as the fraction of paternal mtDNA is boosted by positive selection. The fraction of seeded lineages becomes frozen while intracellular selection within each seeded lineage eventually makes that lineage homoplasmic for the paternal haplotype; the final number of cell lineages homoplasmic for the paternal haplotype is essentially equal to the number of seeded lineages at the time of birth. This observation is illustrated in **Figure S2**. This implies that the level somatic heteroplasmy of the paternal haplotype in the progeny should be approximately equal to the fraction of seeded somatic lineages. Similarly, the paternal inheritance rate (i.e., the proportion of progeny showing measurable paternal heteroplasmy in somatic tissues) should be equal to the fraction of seeded male germline lineages.

The anticipated heteroplasmy levels and inheritance rate of paternal mtDNA, as derived from our simulations (with parameters as described in **Note 7.2**) are presented and compared to the experimentally observed values in **Table1** below. As one can see, the correspondence is fairly good. This good fit is not intended to be used to corroborate our proposed scenario or justify our choice of parameters. Instead this is intended to demonstrate that the ‘impossible’ observations Luo’s 2018 in fact might have resulted from at least one very plausible scenario.

##### Note 7.3.2 Maternal inheritance of the grand-paternal mtDNA

In addition to paternal mtDNA transmission, Luo et al. reports *maternal* transmission of the once paternally inherited haplotype (i.e., haplotype inherited from the male parent at grand-paternal level or even earlier). The authors **incorrectly** call this normal maternal transmission: Unlike normal maternal transmission, the grand-paternal heteroplasmy level in the maternal germline must be diminutively lower than the somatic heteroplasmy of their offspring, because higher maternal germline heteroplasmy would have caused dense seeding of the embryonic cells and resulted in 100% paternal homoplasmy in the offspring due to selection in favor of the grand-paternal haplotype (instead, 22-57% heter*o*plasmy was observed). Our model correctly predicts the observed maternal inheritance rate and heteroplasmy levels of the paternal haplotype (**Note 7.2.4, Table 1**)

**Figure S3.**
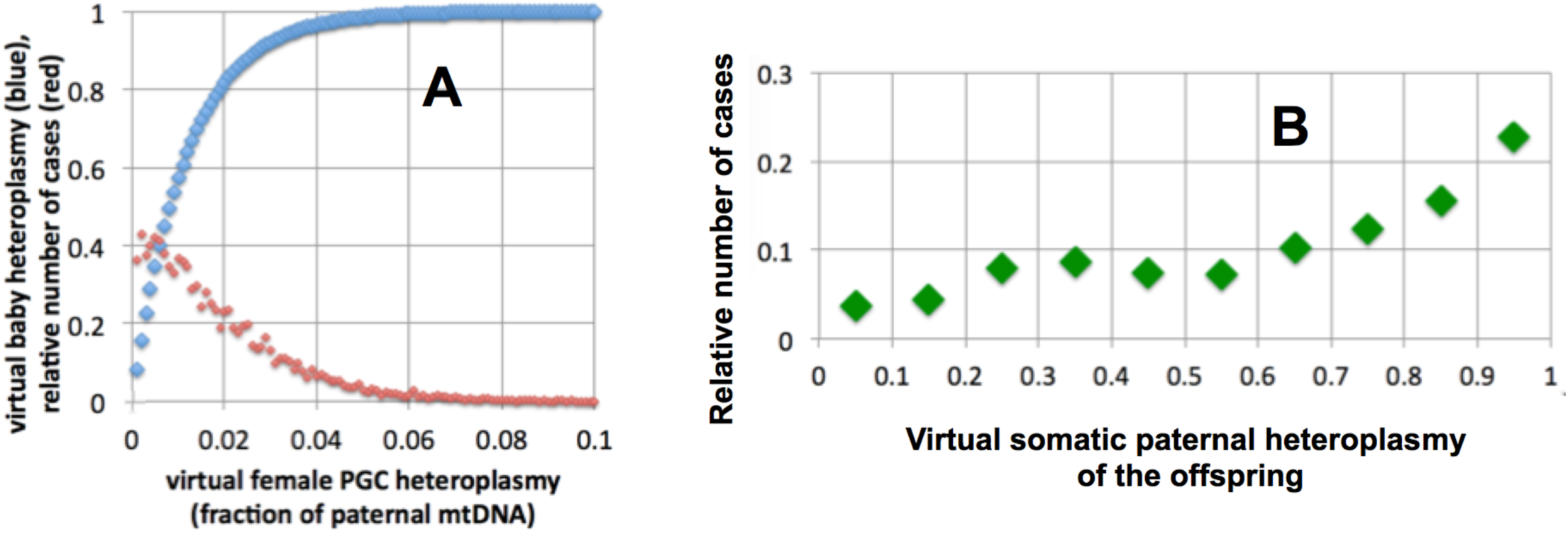
**A**: Red: the distribution of simulated heteroplasmy among virtual PGCs of mothers who inherited a fraction of mtDNA from their fathers. Note the wide range of heteroplasmy levels. Blue: the predicted final *somatic* heteroplasmy levels in virtual offspring resulting from PGCs with the heteroplasmy levels indicated on the X-axis. The saturated appearance of the blue curve reflects the fact that paternal genomes present in PGCs (and in the resulting oocytes) are subject to strong positive enrichment during the development of virtual somatic tissues of the virtual offspring. **B**: The simulated probability distribution of somatic heteroplasmy in the offspring of mothers who had inherited paternal mtDNA. **Note:** zero heteroplasmy cases are not shown (they would have looked like off-scale point at zero).

An explanation for the maternal transmission of the grand-paternal haplotype that has been reported by Luo et al. for 4 cases requires a different approach. Unlike the male germline and somatic cells (**Note 7.3.1**), the female germline is not subject to haplotype selection in keeping with classical results (Jenuth et al., 1997) and it also has very different mtDNA dynamics. The virtual PGCs (primordial germ cells) in our simulations show a wide range of paternal heteroplasmy levels which are transferred almost unchanged to the virtual oocytes derived from these PGCs (red curve in the **Figure S3A** below). Because there is no selection, there is no digitization of the heteroplasmy in individual oocytes (like in spermatozoa). The variable levels of heteroplasmy in the oocytes should be translated into variable levels of somatic heteroplasmy in the offspring (**Fig S3A,** blue curve). The distribution predicts that any level of heteroplasmy is similarly likely to appear in the offspring (**Fig. S3B**), with moderate bias towards the higher levels.

Surprisingly, offspring of the mothers that carry paternal mtDNA (Luo et al.) have a fairly narrow range of heteroplasmy levels: 0.22, 0.22, and 0.29 in a set of 3 siblings from family A and 0.57 in an individual from family C. This distribution appears too narrow to have been *independently* sampled from the distribution shown in Figure S3B, and this was our major concern as this model was being developed. We note however, that sampling may have not been independent. It has been shown (Reizel et al., 2012), that it is not uncommon for a majority of oocytes of the same individual have a common ancestral cell after germline fate has been specified. According to our modeling, heteroplasmy levels in such sibling oocytes should be very uniform. In other words, the three sibling heteroplasmy levels may be so similar because they are closely related, which makes these measurements non-independent and essentially resolves the problem. Assuming that the three siblings in fact represent one ancestral germ cell, we presented their average in table 1.

Of note, our modeling implies that haplotype selection, if it was present in the maternal germline at level similar to that in somatic cells/male germline, would have resulted in very high paternal somatic heteroplasmy in most offspring of mothers inheriting the paternal haplotype. This is because under selection the frequency distribution of PGC/oocyte heteroplasmy (red) in figure S3A will shift to the right, which implies a steep increase in the resulting anticipated heteroplasmy in the offspring (blue curve). This would be in startling contrast with the observed fairly *low* paternal heteroplasmy in these individuals (35% on average, table 1). In other words the independently established lack of selection in the germline is nicely in line with simulations.

##### Note 7.3.3 Note of caution

Here we must once more emphasize that our model contains several inexact parameters and different combinations of these parameters could result in equally realistic predictions. For example, the size of the replicating subpopulation strongly affects the output of the simulations. Smaller subpopulations result in a higher percentage of paternal haplotype in the offspring and an increased inheritance rate (being a part of a smaller replicating subpopulation paternal gives a haplotype a stronger initial boost in relative copy number). Decreasing the size of the replicating subpopulation combined with an even earlier onset of mtDNA replication (which makes sense with a smaller subpopulation) can compensate for a smaller input of mtDNA from the spermatozoid. Neither of these parameters is currently known with certainty (e.g. we do not really know how many mtDNA are delivered by a spermatozoid in the absence of a gatekeeper; 100 copies used in our simulations was no more than a realistic estimation). Thus the results reported herein should not be taken as corroboration of certain values of parameters (such as proving that 10% is the size for replicating subpopulation). Instead the value of these simulations is in demonstrating the importance of these parameters (e.g. without assuming a small replicating mtDNA subpopulation in the early embryo, Luo’s data cannot be explained in a realistic way). In this way, the new cases of paternal inheritance will hopefully serve as catalyst for the studies of mtDNA dynamics in germline and development.

### Note 8. The Vissing case of paternal transmission

The high paternal heteroplasmy in Vissing’s case, i.e., 90% in muscle (Schwartz and Vissing, 2002), appears to be too high for our model. The chances of achieving seeding density of 90% appears low. This difficulty can be resolved, however, if we note that Vissing’s case and Luo’s cases differ in the way the subjects were searched for. Vissing’s patient was discovered as suffering from mtDNA disease, directly related to the pathological and highly detrimental mutation that he had inherited as a part of paternal mtDNA. This means that the search in this case was targeted at high heteroplasmy of the paternal mtDNA, because the mtDNA microdeletion of the type present in this patient would only exert a phenotype when heteroplasmy reaches over ∼90% in muscle. In other words, Vissing’s patient would be invisible to the search had his paternal heteroplasmy not exceeded 90%. Keeping this in mind, we can allow relatively rare circumstances to be used as an explanation for unusually high heteroplasmy levels in this patient (see next paragraph). Note that the search in Luo’s case was *not* for high heteroplasmy, but merely heteroplasmy detectable by sequencing of a blood sample, with the sensitivity of NGS sequencing (∼15-20%).

According to our model, high heteroplasmy can only be achieved via high seeding density of the paternal mtDNA in the cell precursors of the target tissue. There are several ways this can happen as a relatively rare event. One possibility is that the entire tissue, such as muscle in this case, had a relatively recent common ancestor which happened to be highly seeded with paternal mtDNA (such cells do appear at early stages in our models). Indeed, as noted earlier for oocytes, the entire or great majority of cells in a tissue can have a relatively recent common ancestor cell. This has been observed for oocytes and for mesenchymal stem cells (Reizel et al., 2012). The other possibility is that embryo proper in this case happened to be particularly enriched with paternal mtDNA because the early blastomere encompassing the cloud of paternal mtDNA at the point of sperm penetration happened to found a majority of embryo proper cells. Such cases are relatively frequent and have been recorded multiple times in the blastomere lineage tracing experiments (Tabansky et al., 2013). In other words, spatial clustering of the type mentioned in Note 7.2.1, which favored preferential seeding of muscle precursors with paternal haplotype could have resulted in high-density seeding needed to explain high heteroplasmy in Vissing’s patient. Indeed, a little over a 4-fold increase of seeding density (which will happen if paternal mtDNA cluster is entirely encompassed by one blastomere of the 4-cell embryo that then founds the entire embryo proper) is expected to be sufficient to boost heteroplasmy from regular 0.36 to ∼0.90, as observed in Vissing’s patient’s muscle.

Interestingly, all the probands in (Luo et al., 2018) were identified by *suspected mtDNA disease*, as was the original Vissing’s patient. This observation therefore could be somehow related to selective advantage of the paternal haplotype – it’s a known phenomenon that inactivating mtDNA mutations often gain selective intracellular advantage in certain conditions, and there is voluminous literature striving to find the mechanism(s) of this counterintuitive phenomenon (Kowald and Kirkwood, 2018). However, clearly inactivating mtDNA mutations were apparently absent in the reported paternal haplotypes. Alternatively, a mtDNA-disease-like phenotype may be related to the dominant gatekeeper mutations which enabled leakage of sperm mitochondria into the oocyte in these families; such a mutation may have systemic consequences in other aspects of mitochondrial metabolism resulting in the alleged mtDNA-disease-like phenotype in these individuals.

### Note 9. Testable predictions of the model

Single cell analysis should reveal intercellular heteroplasmy and intracellular homoplasmy (mtDNA mosaic). This may be most easily tested in buccal swabs.

### 10. Quality control items

- NGS raw data must be made available to the reader.

- Tests for NUMT contamination must be performed (**Note 1.3**)

- Our own analysis of mitochondrial DNA heteroplasmy in the exome sequencing data from 1339 unrelated individuals revealed three clear instances of multiple heteroplasmies (up to 25% heteroplasmy), potentially originating from mixtures of different mitochondrial haplotypes. However, we failed to confirm these cases of multiple heteroplasmy by conventional Sanger sequencing and the analysis of available family members did not show any evidence for paternal inheritance. Thus, our cases were very likely generated by barcoding errors inherent to modern NGS technology. This proves that bi-paternal mtDNA inheritance is obviously a very rare phenomenon. In light of this rarity an independent confirmation of multiple heteroplasmies in the patients from Luo et al., 2018 by conventional Sanger sequencing would have been desirable, the presented RFLP data appear to us not a sufficient proof to exclude potential methodological problems.

